# Building synthetic chromosomes from natural DNA

**DOI:** 10.1101/2023.05.09.540074

**Authors:** Alessandro L.V. Coradini, Christopher Ne Ville, Zachary A. Krieger, Joshua Roemer, Cara Hull, Shawn Yang, Daniel T. Lusk, Ian M. Ehrenreich

**Author notes:** Correspondence to: Ian M. Ehrenreich, Alessandro L.V. Coradini. Equal contribution authors (Christopher Ne Ville, Zachary A. Krieger).

## Abstract

De novo chromosome synthesis is costly and time-consuming, limiting its use in research and biotechnology. Building synthetic chromosomes from natural components is an unexplored alternative with many potential applications. In this paper, we report CReATiNG (<u>Cl</u>oning, <u>Re</u>programming, and <u>A</u>ssembling <u>Ti</u>led <u>N</u>atural <u>G</u>enomic DNA), a method for constructing synthetic chromosomes from natural components in yeast. CReATiNG entails cloning segments of natural chromosomes and then programmably assembling them into synthetic chromosomes that can replace the native chromosomes in cells. We used CReATiNG to synthetically recombine chromosomes between strains and species, to modify chromosome structure, and to delete many linked, non-adjacent regions totaling 39% of a chromosome. The multiplex deletion experiment revealed that CReATiNG also enables recovery from flaws in synthetic chromosome design via recombination between a synthetic chromosome and its native counterpart. CReATiNG facilitates the application of chromosome synthesis to diverse biological problems.

## Introduction

It is now possible to answer fundamental questions in biology by synthesizing chromosomes (*1, 2*). For example, a longstanding question has been what is the minimal set of genes required to produce a living cell (*3*–*6*)? To answer this question, researchers used design-build-test cycles to synthesize a *Mycoplasma mycoides* chromosome that contains only 473 genes and is still capable of producing a free-living bacterium that replicates on lab timescales (*7*). Among the genes in this minimal set, 83% functioned in the expression and preservation of genetic information, the cell membrane, or cytosolic metabolism, while 17% had unknown functions. Identification of this minimal gene set demonstrates the potential for using chromosome synthesis to understand the defining mechanisms underlying cellular life and its diversity.

To date, synthetic chromosomes have exclusively been generated de novo, through the progressive assembly of small synthetic fragments of DNA into larger molecules via a combination of in vitro and in vivo techniques (*7*–*22*). De novo synthesis is powerful because it allows the complete reprogramming of a chromosome’s sequence and structure. For example, de novo chromosome synthesis was used to generate an *Escherichia coli* strain in which all 18,214 instances of three codons were synonymously reprogrammed, resulting in a strain that utilizes only 61 codons (*18*). In another example, the Sc2.0 community is using de novo chromosome synthesis to generate a strain of the model budding yeast *Saccharomyces cerevisiae* in which all transposable elements have been eliminated and LoxP sites have been incorporated between genes, enabling the generation of random chromosome rearrangements by Cre recombinase (*12*).

The substantial amount of DNA fragment synthesis and assembly involved in de novo chromosome synthesis limits its use in biological research. Reductions in labor and reagent costs are needed to enable biologists to employ chromosome synthesis more widely. Building synthetic chromosomes from cloned segments of natural DNA could be a relatively cheap and fast alternative to de novo chromosome synthesis. Such a method would enable the use of chromosome synthesis in research that does not require complete chromosome reprogramming. For example, projects that could be enabled include mapping the genetic basis of trait differences between individuals and species, probing the structural requirements of chromosomes, and streamlining chromosomes through the systematic elimination of non-essential genetic elements.

In this paper, we introduce CReATiNG (<u>Cl</u>oning, <u>Re</u>programming, and <u>A</u>ssembling <u>Ti</u>led <u>N</u>atural <u>G</u>enomic DNA), a method for building synthetic chromosomes from natural components in *S. cerevisiae*. The first step of CReATiNG is the cloning of natural chromosome segments such that unique adapter sequences are appended to their termini, specifying how these molecules will recombine with each other later when they are assembled. The second step of CReATiNG is co-transforming cloned segments into cells and assembling them by homologous recombination in vivo. Synthetic chromosomes generated with CReATiNG can replace the native chromosomes in cells, making it possible to directly test their phenotypic effects. Here, we describe CReATiNG and demonstrate several of its use cases.

## Results

### A system for in vivo cloning and reprogramming of natural DNA

CReATiNG involves cloning segments of natural chromosomes in yeast donor cells and then programmably assembling these segments into synthetic chromosomes in different recipient cells. To clone a target segment, we co-transform three reagents into donor cells that constitutively express Cas9 (**Figs. 1A** and **B**; **Supplementary Fig. 1**; **Supplementary Table 1**): 1) in vitro transcribed guide RNAs (gRNAs) that direct Cas9 to cut a segment on each side, excising it from a chromosome; 2) a linear Bacterial Artificial Chromosome/Yeast Artificial Chromosome (BAC/YAC) cloning vector flanked by homology to the ends of the segment, enabling integration of the segment into the vector in vivo by homologous recombination; and 3) a repair template comprised of a dominant drug marker flanked with homology arms that allow a cell to reconstitute its broken chromosome by homologous recombination, resulting in replacement of a segment with a marker.

**Figure 1:**
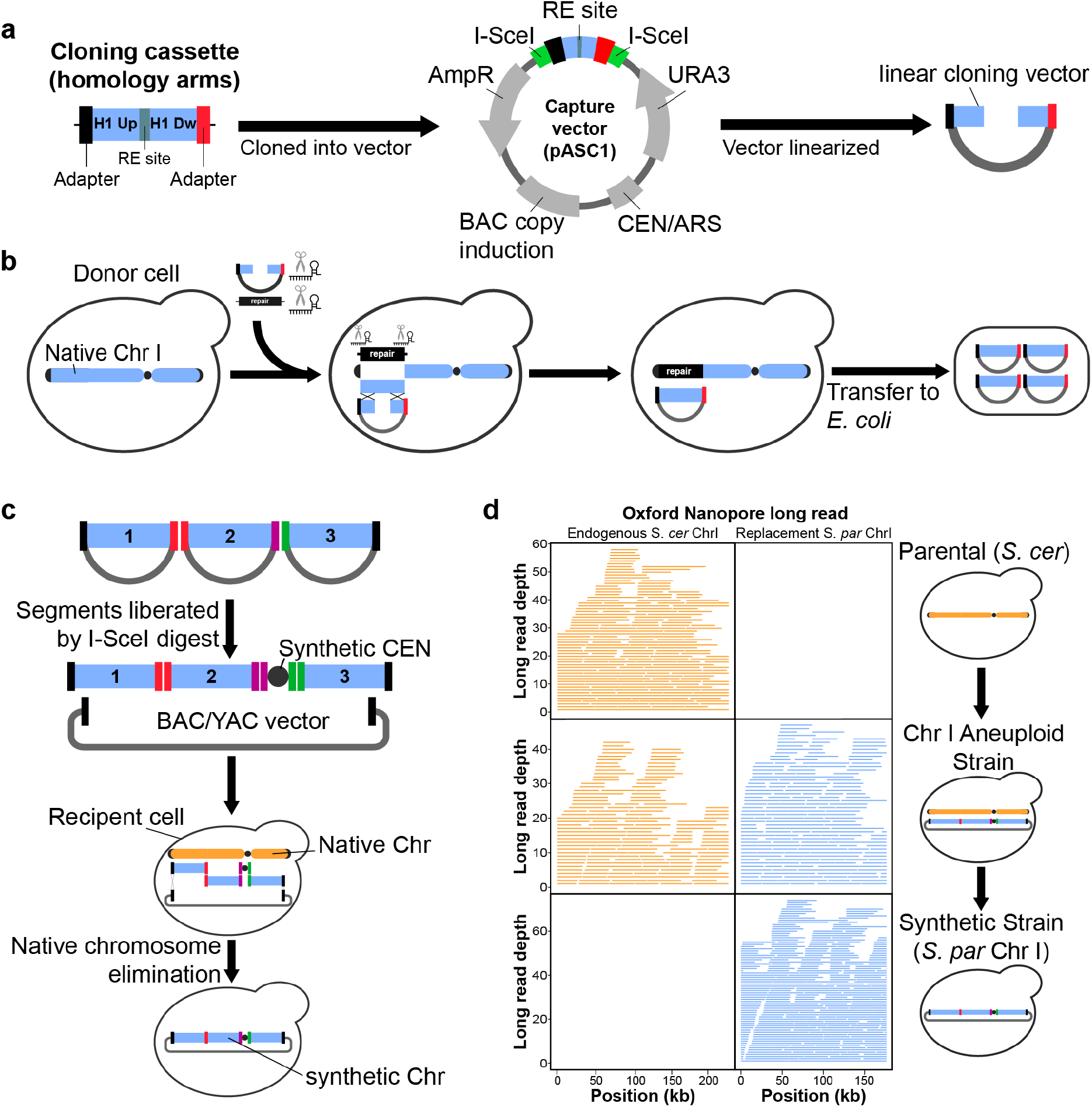
Synthesizing chromosomes from natural components in yeast using CReATiNG. **a**. The Bacterial Artificial Chromosome/Yeast Artificial Chromosome (BAC/YAC) vector used for cloning natural chromosome segments in vivo. Homology may be flanked by sequence adapters that program how a segment will assemble with others in later steps. **b**. A segment is cloned by co-transforming a linearized cloning vector, gRNAs targeting both sides of the segment, and a selectable repair template into a donor cell constitutively expressing Cas9. Cloned segments are then extracted from yeast and transferred to the *E. coli*. **c**. Cloned segments are excised from the vector through restriction digestion with I-SceI. These molecules are then purified and co-transformed into a recipient yeast cell with a centromere cassette and a centromere-free version of the BAC/YAC cloning vector. These molecules are assembled into a synthetic chromosome by homologous recombination in vivo, while the native chromosome is eliminated by centromere destabilization and counterselection. **d**. ONT sequencing confirmed the correct assembly of *S. paradoxus* ChrI (blue) and replacement of the native *S. cerevisiae* ChrI (orange) in BY. The plot shows reads mapped to each chromosome using a reference genome including both BY and *S. paradoxus* ChrI.

Prior to cloning a given target segment, we integrate a segment-specific ∼500 bp cloning cassette that is synthesized de novo into the cloning vector. A cloning cassette contains segment-specific homology arms separated by restriction sites and bounded by I-SceI sites, which are absent from the yeast nuclear genome. Between the I-SceI sites and homology arms, we also include 100 bp DNA sequences (adapters) that are not present in the *S. cerevisiae* genome. These adapters are used to program how segments will assemble later, as different segments with the same adapters will recombine in recipient cells. After transformation with the cloning reagents, donor cells containing successful cloning events can be isolated by selecting for the markers in the cloning vector and the repair template. Cloned segments can then be extracted from yeast donor cells, transformed into *E. coli* for amplification, and extracted from *E. coli* for assembly in yeast recipient cells.

### Initial assembly of a chromosome using CReATiNG

To prototype CReATiNG, we used Chromosome I (ChrI), a 230 kb chromosome containing 117 known or predicted protein-coding genes (*23*). ChrI is the smallest chromosome in the *Saccharomyces* genus and shows synteny between species (*24*). In silico, we divided *S. paradoxus* ChrI into three non-overlapping segments between 51 and 64 kb, which contained the entire chromosome except the centromere, subtelomeres, and telomeres (**Supplementary Fig. 2A**). To assemble segments in their natural order and orientations, we used distinct adapters to specify the junctions between segments 1 and 2, segment 2 and the centromere cassette, and the centromere cassette and segment 3. We excluded subtelomeres from all subsequent work, as they are completely dispensable and highly variable across *Saccharomyces* strains and species (*25*). Telomeres were excluded because they are not amenable to cloning and we assemble chromosomes as circular, rather than linear, molecules.

To enable cloning of the *S. paradoxus* ChrI segments, we generated a Cas9-expressing version of the *S. paradoxus* CBS5829 strain. We then performed three transformations, each targeting a different segment (**Supplementary Table 2 and 3**). For each transformation, we checked five random colonies by amplifying each junction between a cloned segment and the vector (**Supplementary Figs. 2B**-**D**; **Supplementary Table 4**). Based on amplification of both junctions, 14 of 15 (93%) colonies showed successful cloning (**Supplementary Table 5**). After transfer to and amplification in *E. coli*, the three *S. paradoxus* ChrI segments were extracted, liberated from the vector by I-SceI digestion, and purified (**Supplementary Fig. 3A**).

Next, we assembled the segments into a circular chromosome by co-transforming them, the centromere cassette with appropriate adapters, and a centromere-free BAC/YAC into the BY4742 (BY) reference strain of *S. cerevisiae* (**Fig. 1C**). Following transformation, we selected recipient cells in which the five molecules had assembled. Of five colonies checked by PCR of junctions between assembled segments, four (80%) contained the complete assembly (**Supplementary Table 6**). We then performed whole genome Oxford Nanopore Technologies (ONT) sequencing of a single yeast clone and confirmed the presence of two copies of ChrI, one from BY and one from *S. paradoxus* (**Fig. 1D**).

To produce a euploid strain containing only *S. paradoxus* ChrI, we conditionally destabilized and selected against BY ChrI (*12, 26*) (**Supplementary Fig. 3B**). Chromosome elimination involves disrupting centromere function with an inducible *GAL1* promoter and counterselecting a *URA3* marker on the chromosome using 5-FOA. We verified complete elimination of BY ChrI by PCR (**Supplementary Fig. 3C**) and ONT sequencing (**Fig. 1D**). We also used the ONT data to confirm that the euploid strain with synthetic *S. paradoxus* ChrI had the expected sequence and structure genome-wide, with the exception of a single point mutation in a non-functional portion of the centromere cassette (**Supplementary Fig. 4**).

While the remaining work in this manuscript was conducted with circular chromosomes, some chromosome synthesis applications could require linear chromosomes. To confirm that it is possible to convert chromosomes synthesized by CReATiNG from circular to linear forms, we linearized the synthetic *S. paradoxus* ChrI. We used Cas9 to introduce double-strand breaks near each junction between the chromosome and the cloning vector. In addition to gRNAs, we co-transformed repair templates for both chromosome ends, each of which contained a synthetic telomere seed (*27*) and a distinct selectable marker. By selecting for the markers in both repair templates, we obtained cells containing a linear *S. paradoxus* ChrI (**Supplementary Fig. 5**).

### Recombination of chromosomes between strains and species

After confirming that CReATiNG can be used to build synthetic chromosomes that replace the native chromosomes in recipient cells, we explored potential applications. The first application was to synthetically recombine chromosomes between strains and species, which could aid efforts to study the genetic basis of heritable phenotypes. Relative to the crosses conventionally used to generate recombinants, the advantages of CReATiNG are that it does not require mating, meiosis, or natural synteny. Additionally, CReATiNG allows three or more parental chromosomes to recombine in a single assembly. The main constraint of CReATiNG for synthetically recombining chromosomes is that at present it cannot be applied genome-wide.

To use CReATiNG to recombine chromosomes synthetically, we next cloned the three ChrI segments from BY and another *S. cerevisiae* strain, the vineyard isolate RM11-1a (RM). During cloning, we appended the same adapters that were used for *S. paradoxus* segments, making it possible to generate all-possible syntenic combinations of the BY, RM, and *S. paradoxus* segments (**Fig. 2A**). As we had already generated a strain containing an entirely *S. paradoxus* ChrI, we individually assembled the 26 remaining possible chromosomes (**Supplementary Figs. 6 and 7**). These assemblies had efficiencies between 20 and 100% based on PCR examination of five colonies per transformation (**Fig. 2B**; **Supplementary Table 6**). After elimination of native ChrI, we further verified assemblies by ONT sequencing (**Supplementary Fig. 8 and Supplementary Table 7**).

**Figure 2.**
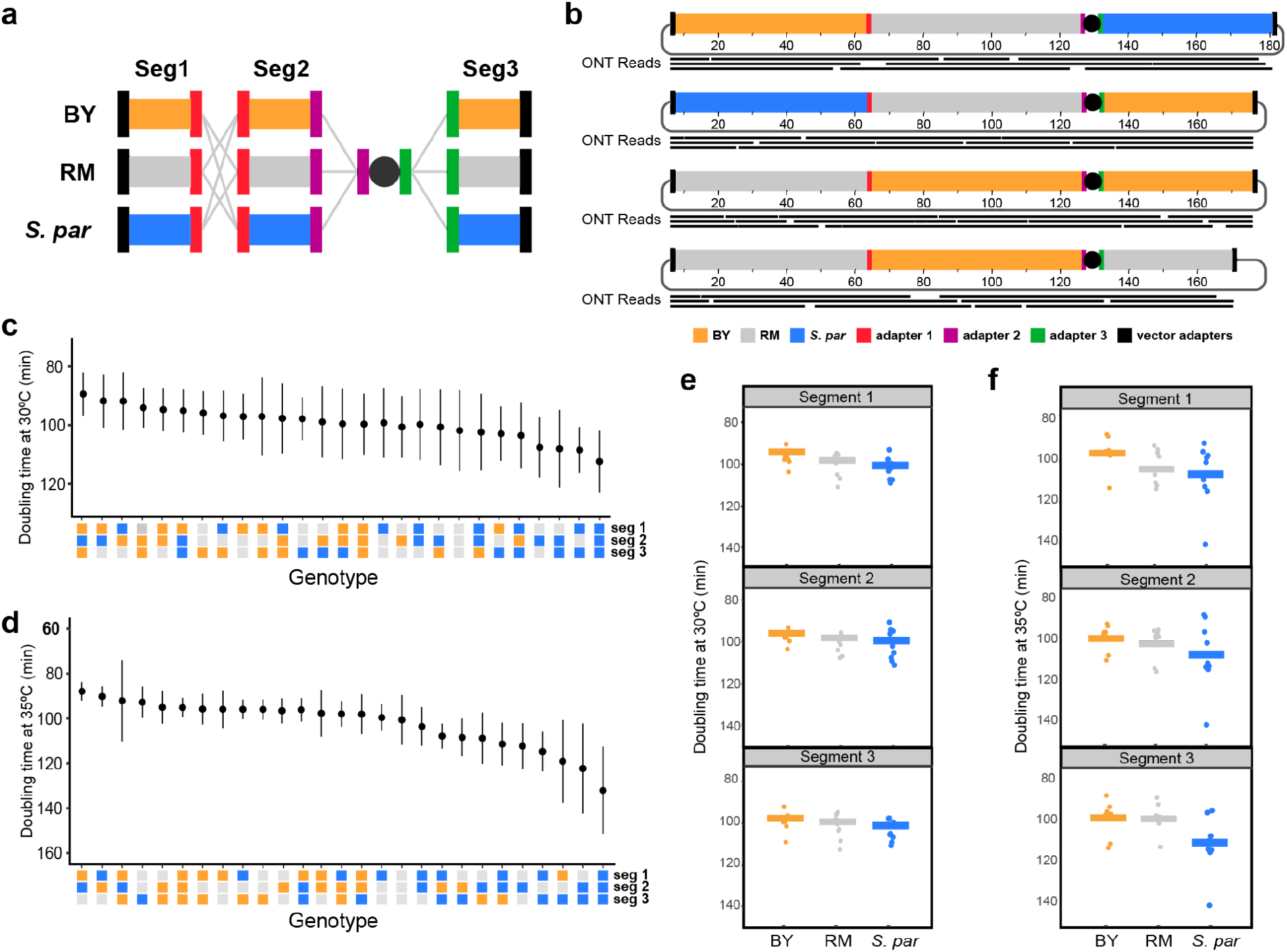
Recombining chromosomes between strains and species using CReATiNG. **a**. Segments 1 through 3 were cloned from two additional strains, the BY (orange) and RM (gray) strains of *S. cerevisiae*. The 27 possible syntenic combinations of the *S. paradoxus* (blue), BY, and RM segments were then assembled in BY recipient cells and the native ChrI was eliminated. **b**. Representative validations of assembled chromosomes by ONT sequencing. In silico designs of each chromosome are shown with a subset of mapped reads plotted below them. **c** and **d**. Euploid strains carrying the synthetic chromosomes were phenotyped for doubling time in rich medium containing glucose at 30ºC and 35ºC, respectively. Means are shown for each strain, with error bars representing one standard deviation around the mean. **e**. The mean effect (horizontal line) of each segment across all genotypes at 30ºC is shown, with dots representing the mean of a genotype across 12 replicates. **f**. The mean effect (horizontal line) of each segment across all genotypes at 35ºC is shown, with dots representing the mean of a genotype across 12 replicates.

Next, we measured the growth rates of these strains with recombinant chromosomes in two conditions–rich liquid medium at 30 and 35°C (**Figs. 2C** and **D**; **Supplementary Table 8**). *S. paradoxus* is known to be more sensitive to high temperature than *S. cerevisiae* (*28*). This phenotypic difference was also present in our donor strains (**Supplementary Fig. 9**). Among the 27 strains with recombinant chromosomes, growth rates varied substantially in both conditions (one-way ANOVAs, p-values ≤ 6.6×10^−11^). In addition, the strain carrying a fully *S. paradoxus* ChrI exhibited the slowest growth at both temperatures, but the difference between it and other strains was more severe at higher temperature. Our results corroborate a recent finding that genetic differences on ChrI contribute to variation in thermotolerance between the two species (*29*).

The 27 strains with recombinant chromosomes also provide an opportunity to measure the individual and combined phenotypic effects of the three ChrI segments. We found that two segments had significant effects on growth at 30°C (one-way ANOVAs, segment 1 p-value = 0.008 and segment 2 p-value = 0.019). However, at 35°C, all three segments had highly significant effects (one-way ANOVAs, p-values ≤ 2.8×10^−5^; **Figs. 2E** and **F**). These results show that all three ChrI segments contribute to growth variation across temperatures, with segment 3 in particular showing a strong interaction with temperature. We also used the data to measure epistasis among the segments and detected significant pairwise and three-way genetic interactions (F tests, p-values ≤ 3.6×10^−4^; **Supplementary Table 9**). Thus, CReATiNG is a useful tool for studying the contribution of genetic factors, including genotype-by-environment interactions and epistasis, to trait differences between strains and species.

### Chromosome restructuring

CReATiNG can also be used to experimentally probe the structural rules underpinning chromosome organization, a topic relevant to genome function and evolution. Recent work suggests yeast can tolerate a diversity of chromosome structures, but most of these studies preserved the order of naturally linked genes (*30*–*33*). CReATiNG makes it possible to restructure the contents of a chromosome in specific non-natural configurations that are programmed using adapters. CReATiNG can be used to synthesize chromosomes with one or more inversions, duplications, deletions, or modifications to gene order.

To demonstrate how CReATiNG can be used in chromosome restructuring, we re-cloned segments 1 through 3 from BY. During this round of cloning, we modified the adapters appended to each segment, making it possible to assemble the segments in all possible orders without inverting any segment. Using these re-cloned segments, we designed five non-natural ChrI structures with the same content but different orders (i.e., 1-3-2, 2-3-1, 2-1-3, 3-1-2, and 3-2-1) (**Fig. 3A**). We produced euploid strains with each non-natural ChrI structure by assembling segments with appropriate adapters and then eliminating the native ChrI. Each assembly was verified by junction PCRs or ONT sequencing (**Supplementary Figure 10**; **Supplementary Table 10**).

While all five strains possessing restructured versions of ChrI were viable, they also showed substantial phenotypic variation. The 2-3-1 configuration exhibited a 7% growth improvement relative to the natural 1-2-3 configuration, which had been generated earlier in the work on synthetic recombinants (**Fig. 3B**; **Supplementary Table 11**). By contrast, the 3-1-2 and 1-3-2 configurations respectively showed growth reductions of 18% and 68% relative to the natural 1-2-3 configuration. Thus, relocating segment 2 to the natural position of segment 3 significantly impedes growth. However, the degree of this impairment also depends on the locations of segments 1 and 3. These findings show how programmably restructuring chromosomes with CReATiNG can be used to identify non-natural chromosome configurations with phenotypic effects.

**Figure 3.**
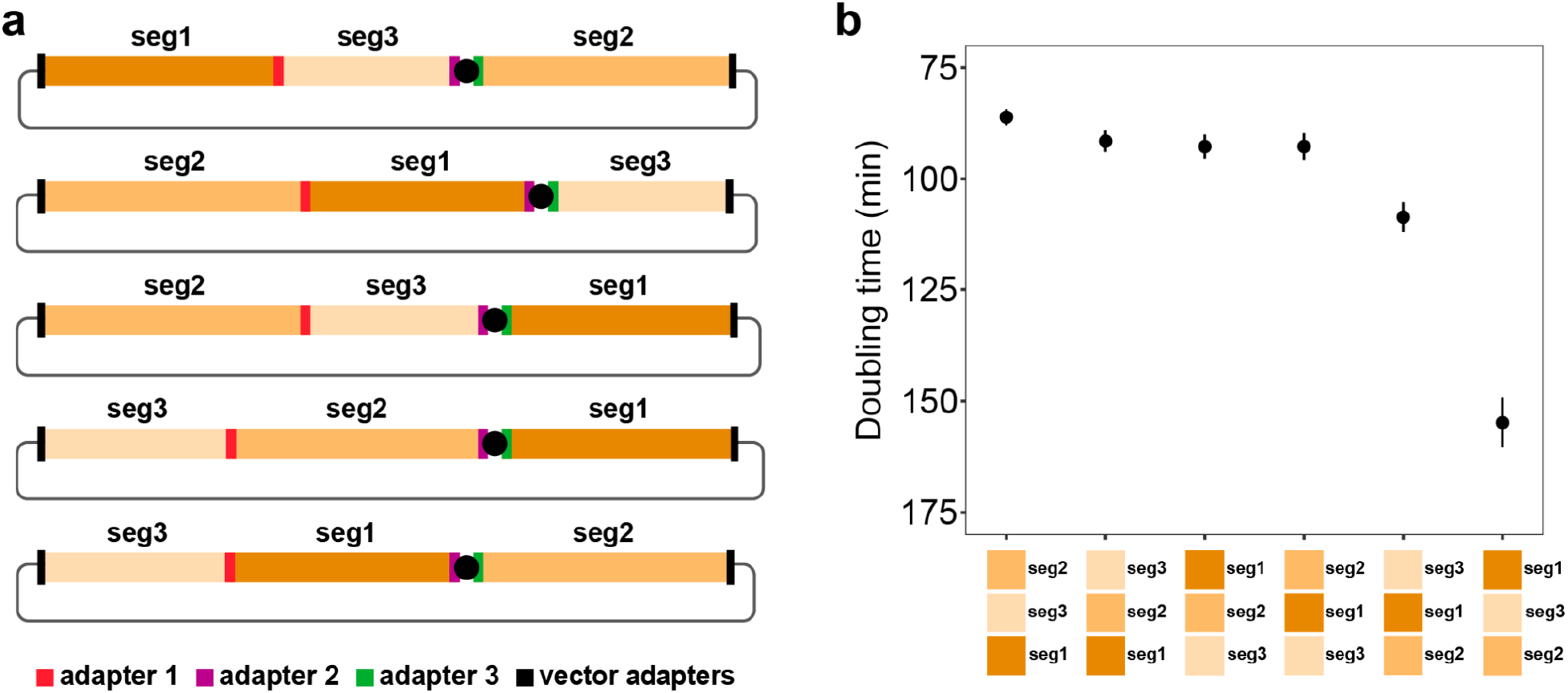
Restructuring ChrI with CReATiNG. **a**. Five possible restructured versions of BY ChrI were created by altering the adapters appended to each segment. In silico designs of each chromosome are shown. PCR of assembly junctions or ONT sequencing was used to confirm that the chromosome assemblies had the correct structure. **b**. Growth analysis of the natural and five non-natural chromosome structures in a rich medium containing glucose at 30ºC. Means are shown for each strain, with error bars representing one standard deviation around the mean.

### Multiplex gene deletion and chromosome streamlining

Another application of CReATiNG is highly multiplexed deletion, a task that remains challenging for conventional genome editing technologies (*34*). Multiplexed deletion could enable the generation of streamlined chromosomes in which many non-essential genetic elements have been eliminated (*35*). Such streamlining can facilitate the production of yeast strains with a substantially reduced gene complement, as well as the reorganization of functionally related genes into modules. CReATiNG simplifies multiplexed deletion: segments of a natural chromosome that should be retained can be cloned and assembled, cleanly deleting all intervening parts of a natural chromosome.

We designed a CReATiNG workflow to delete 10 core chromosome regions from BY ChrI, summing to 30 kb and containing 18 non-adjacent genetic elements in aggregate (12 protein-coding genes, 2 tRNAs, 3 LTRs, and 1 LTR retrotransposon; **Fig. 4A**; **Supplementary Tables 12**-**14**). We chose these elements because they are all annotated as non-essential and no synthetic lethal interactions have been reported among them (*36*). This design required cloning 11 segments, which ranged in size from 3.8 kb to 21 kb (**Supplementary Fig. 11A**; **Supplementary Tables 15** and **16**), and programming them with appropriate adapters for assembly. In BY, the subtelomeres, which we also exclude from synthetic chromosomes, comprise 61.7 kb and 24 protein-coding genes, 1 pseudogene, and 2 LTRs (**Supplementary Table 17**). In total, our multiplex deletion design reduced ChrI by 39.9% (91.7 kb) and eliminated 45 genetic elements.

**Figure 4.**
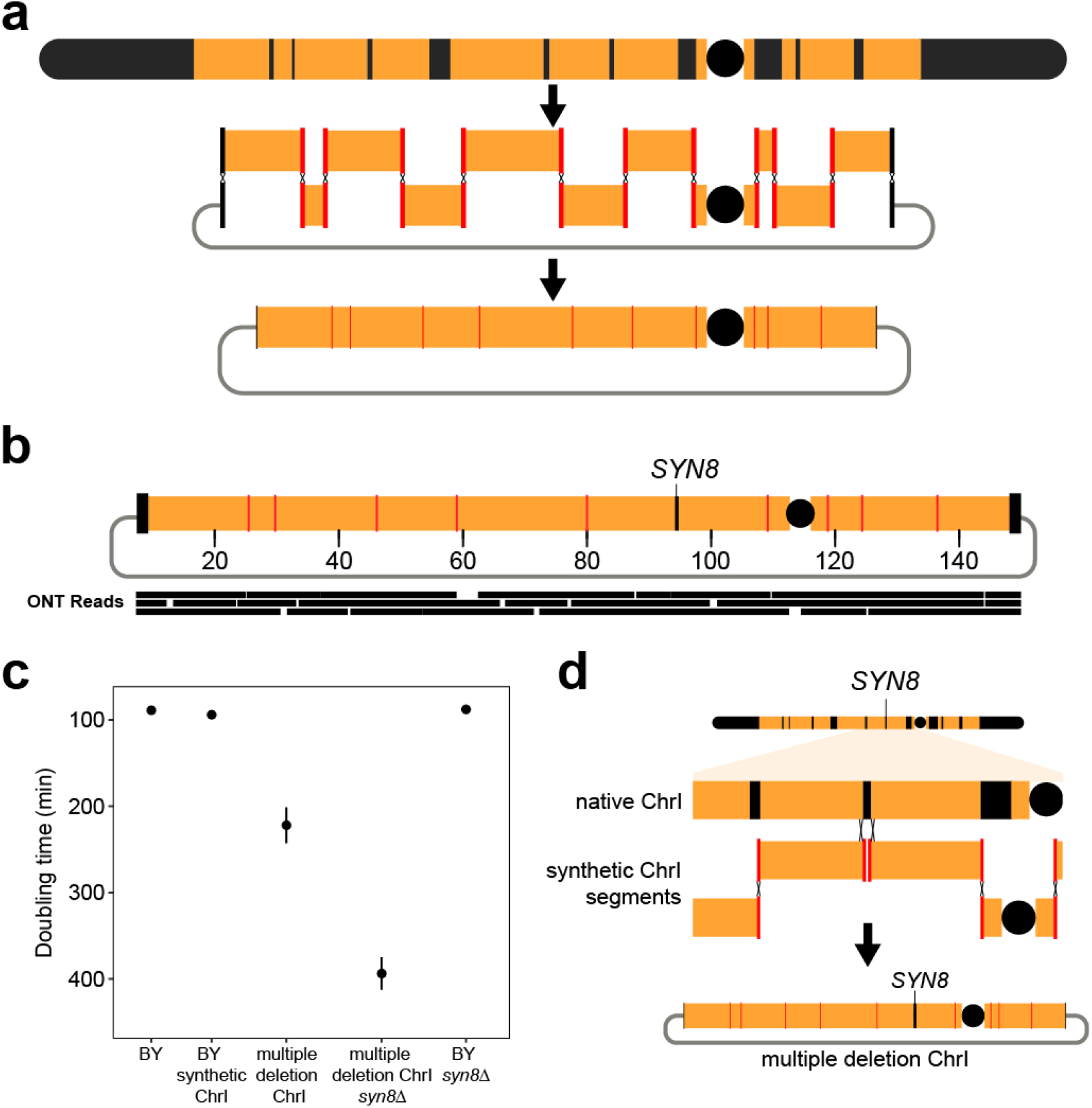
Multiplex gene deletion with CReATiNG. **a**. We attempted to delete 10 non-adjacent regions of the chromosome core and both subtelomeres from BY ChrI, totaling 39.9% of the chromosome. To do this, we cloned 11 segments of the ChrI, assembled them, and performed native chromosome elimination. **b**. ONT sequencing of a colony confirmed results from PCR checks. The colony with the most deletions (nine regions) had the correct structure, but had retained the region containing *SYN8*. An in silico design of the chromosome is shown with a subset of mapped reads plotted below it. **c**. Growth rate analysis of different BY strains, including the unaltered reference strain (BY), a strain carrying a synthetic circular ChrI lacking subtelomeres (BY synthetic ChrI), a synthetic circular ChrI lacking nine core regions and both subtelomeres (multiple deletion ChrI), a synthetic circular ChrI lacking nine core regions, both subtelomeres, and *SYN8* (multiple deletion ChrI *syn8*Δ), and the reference strain with *SYN8* deleted (BY *syn8*Δ). Means are shown for each strain, with error bars representing one standard deviation around the mean. **d**. Recombination between synthetic and native copies of ChrI produced synthetic chromosomes with *SYN8*.

After cloning, we co-transformed the 11 segments into BY along with a centromere cassette and a linearized BAC/YAC vector. *ADE1*, a gene encoding an adenine biosynthesis enzyme whose loss of function causes colonies to accumulate red pigment, was present in one of the regions targeted for deletion. Thus, after native ChrI elimination, we picked 10 red colonies and checked each at all assembly junctions by PCR (**Supplementary Figs. 11B**-**D** and **12**). While none of the red colonies possessed all deletions, a single red colony had nine of the 10 deletions, a finding confirmed by ONT sequencing (**Fig. 4B**). This strain with the multiplex deletion exhibited significantly slower growth than a strain with a synthetic BY ChrI lacking the deletions (212 and 94 minute doubling times, respectively; t-test, p-value = 6.1×10^−11^; **Fig. 4C**).

Of the 10 core chromosome regions targeted for deletion, the only region retained in the colony with nine deletions was between segments 6 and 7. The single gene residing in this region is *SYN8*, which encodes a SNARE protein involved in vesicle fusion with membranes (*36*). Re-examination of the red colonies found that *SYN8* was retained in all 10, suggesting that it genetically interacts with one or more of the other deleted elements. To test for such an interaction, we deleted *SYN8* from the strain with the other deletions. In this context, deleting *SYN8* increased doubling time to 394 minutes, an 86% increase over the multiple deletion strain and a 319% increase over the strain with a synthetic BY ChrI lacking the deletions (**Fig. 4C**; **Supplementary Table 18**). By contrast, *SYN8* deletion had no effect on BY (difference = 1 minute; t-test, p-value = 0.96; **Fig. 4C**; **Supplementary Figure 13**).

Our results imply that multiplex deletion uncovered an unknown genetic interaction between *SYN8* and other elements on ChrI, which converted the normally dispensable *SYN8* into an quasi-essential gene (*7*). Such unknown genetic interactions represent a major obstacle for efforts to streamline chromosomes and genomes, as they will cause strains carrying synthetic chromosomes to show slow growth and poor tractability in the lab. However, the *SYN8* example shows that synthetic chromosomes generated by CReATiNG can overcome such unknown interactions via recombination with native chromosomes prior to native chromosome elimination (**Fig. 4D**).

## Discussion

CReATiNG makes it possible to build synthetic chromosomes with diverse designs using natural components. Because CReATiNG employs cloned segments of natural chromosomes as opposed to small DNA fragments synthesized de novo, it is substantially cheaper and faster than de novo chromosome synthesis. For example, some of the final chromosomes completed for this paper went from in silico design to in vivo testing within a month and cost less than five hundred dollars to produce. Although some synthetic chromosome designs will require complete chromosome reprogramming, which is not possible with CReATiNG, many will not. Indeed, we have shown here that CReATiNG can be used to study a variety of fundamental questions in genetics, genomics, and evolution. Moreover, we unexpectedly found an additional benefit of CReATiNG, which is that it can allow cells to recover from unknown design flaws via recombination between a synthetic chromosome and its native counterpart.

Most of our work in this paper involved simple synthetic chromosome designs in which only three segments were assembled. However, we also demonstrated that CReATiNG can be used to build synthetic chromosomes with complex designs involving ≥10 segments. Using CReATiNG to make synthetic chromosomes with such complex designs could lead to important biological discoveries. For example, here we synthetically recombined chromosomes between strains and species using only two sites. However, this number could be increased, potentially by a large amount, facilitating fine scale genetic mapping of heritable traits within and between species. Similarly, more complex modifications of chromosome structure could be used to identify natural design principles governing chromosome architecture. CReATiNG can also likely be used to delete larger numbers of linked, non-adjacent chromosome regions than explored here, facilitating rapid chromosome streamlining.

In addition, CReATiNG can also be paired with de novo chromosome synthesis to enable projects that might not otherwise be feasible. For example, chromosomes with *Saccharomyces* architecture but sequences from other non-*Saccharomyces* species could be synthesized de novo and then recombined with *Saccharomyces* chromosomes. Such an experiment would facilitate study of the genetic basis of reproductive isolation and trait differences between phylogenetically distant organisms. In addition, CReATiNG and de novo chromosome synthesis could be employed in combination to efficiently relocate genes in the same pathways, complexes, or cellular processes to common genetic modules. Further yet, with some modifications, CReATiNG in yeast may be employed to produce or modify large DNA constructs, including synthetic chromosomes and gene variant libraries, for use in other systems, such as bacteria or mammalian cells. These diverse applications highlight how CReATiNG democratizes the use of chromosome synthesis in biological research.

## Acknowledgments

We thank Oscar Aparicio and Steven Finkel for allowing us to use equipment in their labs and Norman Arnheim, Steven Finkel, Joseph Hale, Julia Schwartzman, and John Tower for feedback on a manuscript draft. This work was supported by startup funds from the University of Southern California, grants 2124400 from the National Science Foundation and R35GM130381 from the National Institutes of Health to I.M.E., as well as an Agilent Postdoctoral Fellowship to A.L.V.C.

## Author Contributions

A.L.V.C. and I.M.E. conceptualized this project. A.L.V.C., C.N.V., Z.A.K., J.R., C.H., S.Y., and D.T.L. performed the experiments. A.L.V.C. and C.N.V. analyzed the data. A.L.V.C., C.N.V. and Z.A.K. generated the figures. A.L.V.C. and I.M.E. wrote and edited the manuscript, with substantial input from C.N.V. and Z.A.K.

